# Assembly and validation of conserved long non-coding RNAs in the ruminant transcriptome

**DOI:** 10.1101/253997

**Authors:** Stephen J. Bush, Charity Muriuki, Mary E. B. McCulloch, Iseabail L. Farquhar, Emily L. Clark, David A. Hume

## Abstract

mRNA-like long non-coding RNAs (lncRNA) are a significant component of mammalian transcriptomes, although most are expressed only at low levels, with high tissue-specificity and/or at specific developmental stages. In many cases, therefore, lncRNA detection by RNA-sequencing (RNA-seq) is compromised by stochastic sampling. To account for this and create a catalogue of ruminant lncRNA, we compared *de novo* assembled lncRNA derived from large RNA-seq datasets in transcriptional atlas projects for sheep and goats with previous lncRNA assembled in cattle and human. Few lncRNA could be reproducibly assembled from a single dataset, even with deep sequencing of the same tissues from multiple animals. Furthermore, there was little sequence overlap between lncRNA assembled from pooled RNA-seq data. We combined positional conservation (synteny) with cross-species mapping of candidate lncRNA to identify a consensus set of ruminant lncRNA and then used the RNA-seq data to demonstrate detectable and reproducible expression in each species. The majority of lncRNA were encoded by single exons, and expressed at < 1 TPM. In sheep, 20-30% of lncRNA had expression profiles significantly correlated with neighbouring protein-coding genes, suggesting association with enhancers. Alongside substantially expanding the ruminant lncRNA repertoire, the outcomes of our analysis demonstrate that stochastic sampling can be partly overcome by combining RNA-seq datasets from related species. This has practical implications for the future discovery of lncRNA in other species.

## Introduction

Mammalian transcriptomes include many long non-coding RNAs (lncRNAs), a collective term for transcripts of > 200 nucleotides that resemble mRNAs (many being 3’ polyadenylated, 5’ capped and spliced) but do not encode a protein product [1]. Proposed functional roles of lncRNAs include transcriptional regulation, epigenetic regulation, intracellular trafficking and chromatin remodelling (see reviews [2-9]). Some view lncRNAs as transcriptional noise [10, 11]. Full length lncRNAs are difficult to assemble: many are expressed at low levels [12], with high tissue-specificity [13, 14], at specific developmental time points (e.g. [15-17]), and with few signs of selective constraint [18, 19]. Many are also expressed transiently, and so may be partly degraded by the exosome complex [20].

The initial recognition of lncRNAs as widespread and *bona fide* outputs of mammalian transcription was based upon the isolation and sequencing of large numbers of mouse and human full-length cDNAs [21-23], many of which were experimentally validated [24] and shown to participate in sense-antisense pairs [25]. They were captured in significant numbers because the cDNA libraries were subtracted to remove abundant transcripts. More recent studies have used RNA-sequencing (RNA-seq) to assemble larger catalogues of lncRNAs [26]. Because of the power-law relationship of individual transcript abundance in mammalian transcriptomes [27], unless sequencing is carried out at massive depth, the exons of lowly-abundant transcripts (such as lncRNAs) are subject to stochastic sampling and are detected inconsistently between technical replicates of the same sample [28]. RNA-seq is also a relatively inaccurate means of reconstructing the 5’ ends of transcripts [29]. To overcome this constraint, the FANTOM Consortium supplemented RNA-seq with Cap Analysis of Gene Expression (CAGE) data, characterising – in humans – a 5’-complete lncRNA transcriptome [30].

RNA-seq libraries from multiple tissues, cell types and developmental stages are commonly pooled to maximise the number of lncRNA gene models assembled. Genome-wide surveys have expanded the lncRNA repertoire of livestock species such as cattle (18 tissues, sequenced at approx. 40-100 million reads each) [31], pig (10 tissues, sequenced at approx. 6-40 million reads each) [32], and horse (8 tissues, sequenced at approx. 20-200 million reads each) [33], complementing tissue-specific lncRNA catalogues of, for example, cattle muscle [34, 35] and skin [36], and pig adipose [37, 38], liver [39] and testis [40].

The low level of lncRNA conservation (at some loci, it appears that only the act of transcription, rather than the transcript sequence itself, is functionally relevant [41]) reduces the utility of comparative analysis of the large RNA-seq datasets available from human [30, 42] and mouse [43]. Amongst 200 human and mouse lncRNAs, each characteristic of specific immune cell types, there was <1% sequence conservation [44].

Here we focus on more closely related species. We have generated atlases of gene expression for the domestic sheep, *Ovis aries* [45], and the goat, *Capra hircus* (manuscript in preparation). As the two species are closely related (sharing a common ancestor < 10mya [46]) and their respective RNA-seq datasets contain many of the same tissues, it is possible to use data from one species to infer the presence of lncRNAs in the other. Cattle and humans are more distantly related to small ruminants, but nevertheless are substantially more similar than mice. We extend our approach by utilising existing human and cattle lncRNA datasets to identify a consensus ruminant lncRNA transcriptome, and use the sheep transcriptional atlas to confirm that candidate lncRNA identified by cross-species inference are reproducibly expressed. The lncRNA catalogues we have generated in the sheep and goat are of interest in themselves [47] and contribute valuable information to the Functional Annotation of Animal Genomes (FAANG) project [48, 49].

## Results and Discussion

### Identifying lncRNAs in the sheep and goat transcriptomes

We have previously created an expression atlas for the domestic sheep [45], using both polyadenylated and rRNA-depleted RNA-seq data collected primarily from three male and three female adult Texel x Scottish Blackface (TxBF) sheep at two years of age: 441 RNA-seq libraries in total, comprising 5 cell types and multiple tissues spanning all major organ systems and several developmental stages, from embryonic to adult. To complement this dataset, we also created a smaller-scale expression atlas – of 54 mRNA-seq libraries – from 6 day old crossbred goats, which will be the subject of a dedicated analysis. For both species, each RNA-seq library was aligned against its reference genome (Oar v3.1 and ARS1, for sheep and goat, respectively) using HISAT2 [50], with transcripts assembled using StringTie [51]. This pipeline produced a non-redundant set of *de novo* gene and transcript models, as previously described [45], and expanded the set of transcripts in each reference genome to include *ab initio* lncRNA predictions and novel protein-coding genes. As the primary purpose of the sheep expression atlas was to improve the functional characterisation of the protein-coding transcriptome, the novel sheep protein-coding transcript models generated by this pipeline have been previously discussed [45] (novel protein-coding transcripts for goats will be discussed in a dedicated analysis of the protein-coding goat transcriptome).

Using similar filter criteria to a previous study [52], the *de novo* gene models were parsed to create longlists of 30,677 (sheep) and 7671 (goat) candidate lncRNAs, each of which was >= 200bp and was not associated, on the same strand, with a known protein-coding locus. The 4-fold difference in the length of each longlist can be attributed to the relative size of each dataset. The sheep atlas contains 8 times as many RNA-seq libraries, spans multiple developmental stages (from embryonic to adult), and has a subset of its samples specifically prepared to ensure the comprehensive capture of ncRNAs – unlike any sample in the goat dataset, this subset is sequenced at a 4-fold higher depth (>100 million reads, rather than >25 million reads) using a total RNA-seq, rather than mRNA-seq, protocol.

Each model on both longlists was assessed for coding potential using the classification tools CPC [53], CPAT [54] and PLEK [55], alongside homology searches of its longest ORF – with blastp [56] and HMMER [57] – to known protein and domain sequences (within the Swiss-Prot [58, 59] and Pfam-A [60] databases, respectively). Those gene models classified as non-coding by CPC, CPAT and PLEK, and having no detectable blastp and HMMER hits, are considered novel lncRNAs.

This pipeline creates shortlists of 12,296 (sheep) and 2657 (goat) lncRNAs (Tables S1 and S2, respectively), representing approximately 40% (sheep) and 35% (goat) of the gene models on each longlist. The mean gene length is similar in both shortlists – 6.7kb (sheep) and 8.8kb (goat) – as is summed exon length, averaging 1.2kb in each species.

Consistent with previous analysis in several other species [31, 61], 6956 (57%) of the sheep lncRNAs, and 1284 (48%) of the goat, were single-exonic. For sheep, the shortlist contains 11,646 previously unknown lncRNA models and provides additional evidence for 650 existing Oar v3.1 lncRNA models (Table S1). A small proportion of longlisted gene models were considered non-coding by at least one of CPC, CPAT or PLEK, but nevertheless showed some degree of sequence homology to either a known protein or protein domain: for sheep, 226 (including 13 existing Oar v3.1 models) (Table S3), and for goats, 153 (Table S4). The number of novel lncRNAs identified is also given per chromosome (Tables S5 (sheep) and S6 (goat)) and per type (Tables S7 (sheep) and S8 (goat)), the majority of which – in both species – are found in intergenic regions, 10-100kb from the nearest gene. Overall, these lncRNA models increase the number of possible genes in the reference annotation by approximately 30% (sheep) and 12% (goat).

### The sets of ab initio sheep and goat lncRNAs only minimally overlap at the sequence level

Even with full length cDNA sequences, comparative analysis revealed that only 27% of the lncRNAs identified in human had mouse counterparts [23]. When comparing the sets of sheep and goat lncRNAs, few predicted transcripts – in either species – show sequence-level similarity either to each other or to other closely or distantly related species (cattle and human, respectively, which shared a common ancestor with sheep and goats approx. 25 and 95mya [46]). Of the 12,296 shortlisted sheep lncRNAs, less than half (n = 5139, i.e. 42%) had any detectable pairwise alignment – of any quality and of any length – to either the shortlisted goat lncRNAs, a set of 9778 cattle lncRNAs from a previous study [31] or two sets of human lncRNAs (Figure 1 and Table S9). In only a small proportion of these alignments can there be high confidence: that is, the alignment has a % identity >= 50% within an alignment >= 50% the length of the target sequence. Of the 5139 sheep lncRNAs that could be aligned to any species, only 293 (5.7%) could be aligned with high confidence to goat and 265 (5.2%) to cattle transcripts. Similarly, of the sheep lncRNAs that could be aligned to either of two human lncRNA databases – NONCODE [62] and lncRNAdb [63] – 68 (1.6% of the total alignable lncRNAs) aligned with high confidence to the NONCODE database, and none to the lncRNAdb. Similar findings are observed with the 2657 shortlisted goat lncRNAs: 1343 (50.5%) had a detectable pairwise alignment, of any quality, to either set of sheep, cattle or human lncRNAs. However, of these 1343 lncRNAs, only 113 (8.4%) aligned with high confidence to sheep, 88 (6.6%) to cattle, 55 (4.1%) to the human NONCODE database, and 1 (0.1%) to the human lncRNAdb database (Figure 1 and Table S10). These observations allow for two possibilities. Firstly, lncRNAs may, in general, be poorly conserved at the sequence level, consistent with previous findings [18, 19] and the observation that only 6% of the sheep/goat alignments have >50% reciprocal identity.

**Figure 1.**
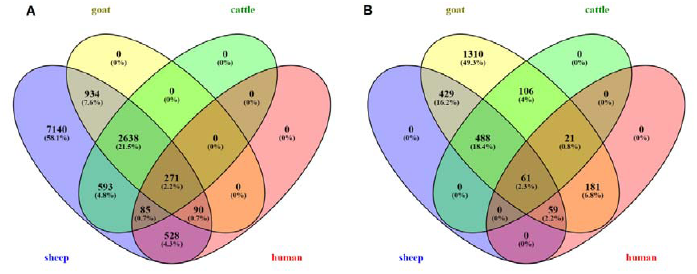
Minimal overlap of lncRNAs at the sequence level. Venn diagrams show the number of sheep (A) or goat (B) lncRNAs that can be aligned – with an alignment of any length or quality – to either shortlist of goat (A) or sheep (B) lncRNAs, and to sets of cattle and human lncRNAs from previous studies. The majority (58% of sheep lncRNAs, and 49% of goat lncRNAs) have no associated alignment. Alignments are detailed in Tables S9 (sheep) and S10 (goat).

However, an alternative is that despite the apparent depth of coverage, we have only assembled a subset of the total lncRNA transcriptome in each species.

### lncRNAs not captured by the RNA-seq libraries of one species can be found using data from a related species

A reasonable *a priori* prediction is that lncRNAs – if functionally relevant – are most likely to share expression in a closely related species. Whereas human and mouse lncRNAs identified as full length cDNAs were generally less conserved between species than the 5’ and 3’UTRs of protein-coding transcripts, their promoters were more highly conserved than those of protein-coding transcripts, some extending as far as chicken [43, 64]. These findings suggested that the large majority of lncRNAs that were analyzed displayed positional conservation across species. Accordingly, rather than comparing the similarity of two sets of lncRNA transcripts, we mapped the lncRNAs assembled in one species (e.g. sheep) to the genome of another (e.g. goat), deriving confidence in the mapping location from synteny. For each of the pairwise sheep/cattle, sheep/goat, cattle/goat, sheep/human, goat/human, and cattle/human comparisons, we identified sets of syntenic blocks: regions in the genome where gene order is conserved both up‐ and downstream of a focal gene (see Table 1 and Methods). In the sheep/cattle comparison, approximately 5% of the syntenic blocks contain at least one lncRNA with a relative position conserved in both species, either upstream (n=139 lncRNAs) or downstream (n=141) of the central gene in each block (Table S11). In the sheep/goat and cattle/goat comparisons, respectively, approximately 2 and 3% of the syntenic blocks contain a lncRNA (for sheep/goat, n=42 upstream, 40 downstream; for cattle/goat, 86 upstream, 83 downstream) (Tables S12 and S13, respectively). With increased species divergence, far fewer lncRNAs (<1%) have relative positions conserved in either the upstream or downstream positions of the sheep/human, goat/human and cattle/human syntenic blocks (Tables S14, S15 and S16, respectively). These comparatively small proportions highlight the minimal overlap between each set of assembled transcripts, consistent with stochastic assembly – lncRNAs expected to be present in a particular location are captured in only one species, not both. As such, very few lncRNAs in either of the sheep, goat and cattle subsets have evidence of both shared sequence homology and conserved synteny. When comparing sheep and cattle, 16 unique lncRNAs have high-confidence pairwise alignments within a region of conserved synteny, and when comparing sheep and goat, 6 (Table S17).

**Table 1.** Comparatively few lncRNAs appear positionally conserved, suggesting minimal overlap between each species’ set of assembled transcripts. This suggests that those lncRNAs expected to be found at a given genomic location are captured in only one species, not both, consistent with the stochastic sampling of lncRNAs by RNA-seq libraries.

| Species 1 | Species 2 | No. of syntenic blocks (i.e. three conserved consecutive genes) | No. of unique protein-coding genes in the set of syntenic blocks | Total no. of positionally conserved lncRNAs in the set of syntenic blocks (in either the up- or downstream position) | % of syntenic blocks with at least one positionally conserved lncRNA |
| --- | --- | --- | --- | --- | --- |
| sheep | cattle | 2927 | 5601 | 280 | 9.57 |
| sheep | goat | 2038 | 3883 | 82 | 4.02 |
| sheep | human | 380 | 930 | 8 | 2.11 |
| goat | cattle | 2982 | 5258 | 169 | 5.67 |
| goat | human | 527 | 1262 | 2 | 0.38 |
| cattle | human | 443 | 1063 | 5 | 1.13 |

In most of the syntenic blocks examined, if a lncRNA was detected in one location in one species (either up‐ or downstream of a focal gene), no corresponding assembled lncRNA was annotated in the comparison species, even though both species sequenced a similar range of tissues. For example, of the 2927 syntenic blocks in the sheep/cattle comparison, 347 (12%) of the sheep blocks, and 506 (17%) of the cattle blocks, contain a lncRNA in the ‘upstream’ position (that is, between genes 1 and 2), with little overlap between the two species: in only 139 blocks (5%) is a lncRNA present in this position in both species (Table S11). Similar results are found if considering the ‘downstream’ position, as well as the sheep/goat, goat/cattle, sheep/human, goat/human and cattle/human comparisons: approximately 2-5 times as many lncRNAs are found in either of the two species than are found in both (Tables S11, S12, S13, S14, S15 and S16).

Each set of syntenic blocks, by definition, represents a set of conserved intergenic regions. Given that the majority of lncRNAs are intergenic (Tables S7 and S8), these regions are reasonable locations for directly mapping candidate transcripts (strictly speaking, concatenated exon sequences) to the genome. For the syntenic blocks in each species comparison, we made global alignments of the lncRNAs in species *x* to the intergenic region of species *y*, and vice versa (see Methods). Retaining only those alignments in which the lncRNA can match the intergenic region with 20 or more consecutive residues (the majority of these alignments in any case have >= 75% identity across their entire length), we predicted 1077 additional lncRNAs in cattle, 1401 in sheep, and 1735 in goat, although only 44 in human (Table 2 and Table S18). That comparatively few ruminant lncRNAs are recognisable at the sequence level in humans (and vice versa) is consistent with the rapid turnover of the lncRNA repertoire between species [65]. In the case of the goat, the number of new lncRNAs predicted by this approach is > 50% the number captured (and shortlisted) using goat-specific RNA-seq (Figure 2). This suggests that for the purposes of lncRNA detection, datasets from related species can help overcome limitations of sequencing breadth and depth. This is even apparent with comparatively large datasets – the sheep RNA-seq, for instance, spans more tissues and developmental stages than goat, but in absolute terms, it still fails to generate assemblies of many lncRNAs.

**Figure 2.**
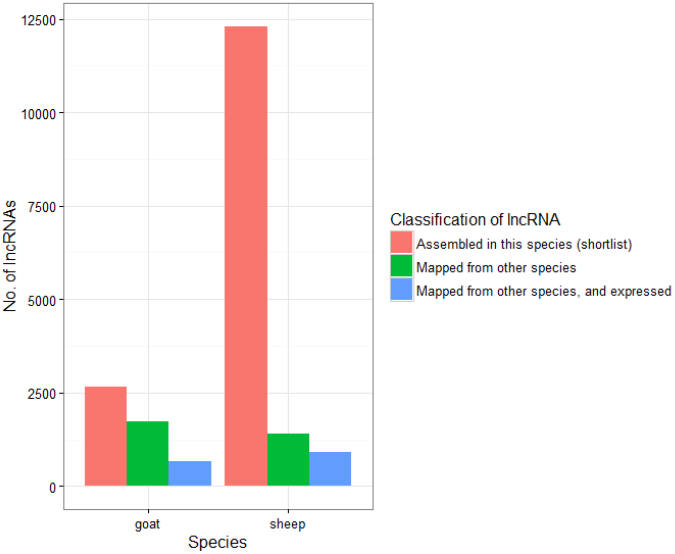
The stochastic detection and assembly of lncRNAs by RNA-seq libraries – a consequence of limitations in sequencing breadth and depth – suggests that for a given species, only a subset of the total lncRNRA transcriptome is likely to be captured. Nevertheless, the number of candidate lncRNAs for that species can be increased if directly mapping, to a positionally conserved region of the genome, the lncRNAs from either a related (sheep, goat, cattle) or more distant (human) species. Many of these mapped lncRNAs (which could not be completely reconstructed with the RNA-seq libraries of that species) are nevertheless detectably expressed.

**Table 2.**
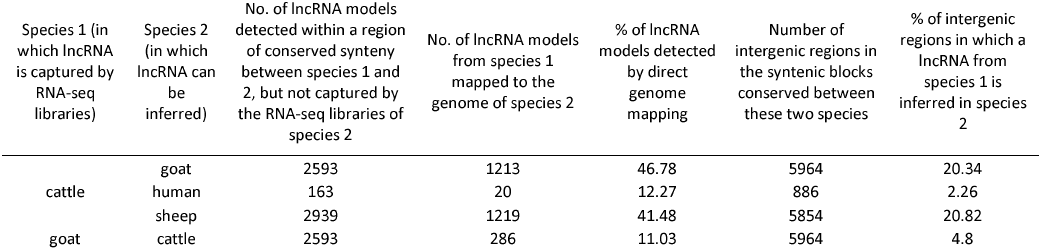
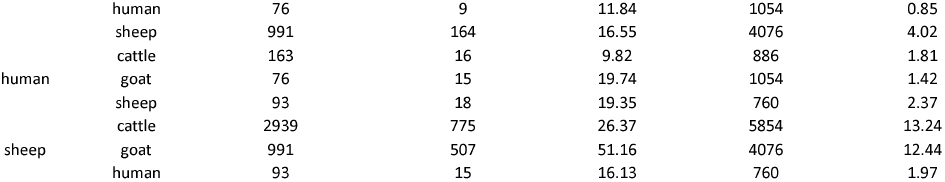
lncRNA transcripts assembled using the RNA-seq libraries of only one species can in many cases be directly mapped to the genome of nother species, assuming the lncRNA is located within a region of conserved synteny.

### Many of the sheep lncRNAs inferred by synteny – which could not be fully assembled from the RNA-seq reads – are nevertheless detectably expressed

To determine the expression level of the sheep lncRNAs, we utilised a subset of 71 high-depth (>100 million reads) RNA-seq libraries from the sheep expression atlas [45]. This subset constitutes a set of 11 transcriptionally rich tissues (bicep muscle, hippocampus, ileum, kidney medulla, left ventricle, liver, ovary, reticulum, spleen, testes, thymus), plus one cell type in two conditions (bone marrow derived macrophages, unstimulated and 7 hours after simulation with lipopolysaccharide), each of which was sequenced in up to 6 individuals (where possible, 3 adult males and 3 adult females).

For each sample, expression was quantified – as transcripts per million (TPM) – using the quantification tool Kallisto [66]. Kallisto quantifies expression by matching k-mers from the RNA-seq reads to a pre-built index of k-mers, derived from a set of reference transcripts. For sheep, we supplemented the complete set of Oar v3.1 reference transcripts (n=28,828 transcripts, representing 26,764 genes) both with the shortlist of 11,646 novel lncRNAs (each of which is a single-transcript gene model) (Table S1), and those lncRNAs assembled from either human, goat and cattle (respectively, 18, 164 and 1219 lncRNAs), whose presence was predicted in sheep by mapping the transcript to a conserved genomic region (Table S18).

Of these 13,047 novel lncRNAs, 8826 were detected at a level of TPM > 1 in at least one of the 71 adult samples, including 14 of the human transcripts (78%), 128 of the goat transcripts (78%), and 772 of the cattle transcripts (63%) (Table S19). At a depth of coverage of 100 million reads, we would expect to detect transcripts reproducibly at between 0.01 and 0.1 TPM if they are expressed in all libraries derived from the same tissue/cell type. Indeed, of the 13,047 total novel lncRNAs, 5353 (41%) were detected with at least one paired-end read in all 6 replicates of the tissue in which it is most highly expressed (Table S19). Those lncRNAs derived from goat and cattle transcripts are similarly reproducible: 83 (51%) of the goat transcripts were detected with at least one paired-end read in all 6 replicates of its most expressed tissue, as were 570 (47%) of the cattle transcripts, and 7 (39%) of the human transcripts (Table S19).

By extension, we can consider sheep, cattle and human lncRNA to be goat lncRNA, and create a Kallisto index containing candidate lncRNAs extracted from the goat genome after mapping sheep and cattle transcripts. Using such a Kallisto index (which contains the 2657 shortlisted goat lncRNAs (Table S2), 507 sheep lncRNAs, 1213 cattle lncRNAs, and 15 human lncRNAs), 1478 (34%) of a total set of 4392 candidate goat lncRNAs were reproducibly detected (> 0.01 TPM) in all 4 of the goats sampled (Table S20). Hence, data from the sheep expression atlas can be used to provide additional functional annotation of the goat genome, despite the much lower number of tissue samples relative to sheep.

In general, lncRNA expression is low: 12,325 sheep lncRNAs (94% of the total) have a mean TPM, across all 71 samples, below 10. The mean and median maximum TPM for each lncRNA across the total sheep dataset was 18.4 and 2.2 TPM, respectively (Table S19). Other reports have described pervasive, but low-level, mammalian lncRNA transcription [12], and – given the mean TPM exceeds the median – a high degree of lncRNA tissue-specificity [67-69]. Indeed, for those lncRNAs detected at > 1 TPM, the average value of *tau* – a scalar measure of expression breadth bound between 0 (for housekeeping genes) and 1 (for genes expressed in one sample only) [70] (see Methods) – is 0.66. Although most of the lncRNAs (n = 4972, 64% of the 7809 lncRNAs with average TPM > 1 in at least one tissue) have idiosyncratic ‘mixed expression’ profiles (see Methods), 1339 lncRNAs (17%) are nevertheless detected at an average TPM > 1 in all 13 tissues (Table S19). Many are enriched in specific tissues, with 904 (12%) lncRNAs exhibiting a testes-specific expression pattern, consistent with a previous study identifying numerous lncRNAs involved in ovine testicular development and spermatogenesis [71].

### Few lncRNAs are fully captured by biological replicates of the same RNA-seq library

In the largest assembly of predicted lncRNAs, from humans, the transfrags (transcript fragments) assembled from 7256 RNA-seq libraries were consolidated into 58,648 candidate lncRNAs [72]. Before assembling transfrags, machine learning methods were employed to filter, from each library, any library-specific background noise (genomic DNA contamination and incompletely processed RNA). Filtered libraries were then merged before assembling the final gene models, in effect pooling together transfrags (which may be partial or full-length transcripts) from all possible libraries. Consequently, a given set of transfrags can be assembled into a consensus transcript for a lncRNA, but that consensus transcript might not actually exist in any one cellular source. The only unequivocal means to confirm the full length expression would be to clone the full length cDNA. However, additional confidence can be obtained by increasing the depth of coverage in the same tissue/cell type in a technical replicate. In the sheep expression atlas, 31 diverse tissues/cell types were sampled in each of 6 individual adults (3 females, 3 males, all unrelated virgin animals approximately 2 years of age). By taking a subset of 31 common tissues per individual, each of the 6 adults was represented by ∼0.75 billion reads.

In a typical lncRNA assembly pipeline, read alignments from all individuals are merged, to maximise the number of candidate gene models (using, for instance, StringTie ‐‐merge; see Methods). With *n* = 6 adults (and ∼0.75 billion reads per adult), there are 2*^n^*-1 = 63 possible combinations of data for which GTFs can be made with StringTie ‐‐merge. The reproducibility of each shortlisted lncRNA, in terms of the number of GTFs it is reconstructed in, is shown in Table S21. The GTFs themselves are available as Dataset S1 (available via the University of Edinburgh DataShare portal; http://dx.doi.org/10.7488/ds/2284).

Only 812 of the 12,296 sheep lncRNRAs (6.6%) could be fully reconstructed by any of the 63 GTF combinations (Table S21). One caveat in this assessment is that these sheep libraries are exclusively from adults. Many of the 12,296 lncRNA models may instead be expressed during embryonic development. There is evidence of extensive embryonic lncRNA expression in human [15, 73] and mouse [16, 74]. The lack of embryonic tissues could also explain why fewer lncRNAs were assembled in goat. Nevertheless, when considering all 429 RNA-seq libraries in the sheep expression atlas (i.e. including non-adult samples), there are only, on average, 29 libraries (7%) in which any individual lncRNA can be fully reconstructed (Figure 3 and Table S22).

**Figure 3.**
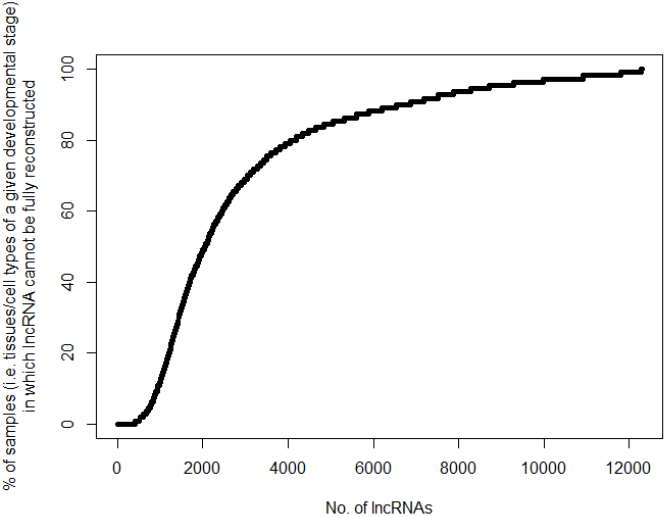
Proportion of samples in the sheep expression atlas for which a candidate lncRNA (n = 12,296) cannot be fully reconstructed. The atlas comprises 429 RNA-seq libraries, representing 110 distinct samples; that is, each sample is a tissue/cell type at a given developmental stage, with up to 6 replicates per sample. 22 candidate lncRNAs cannot be reconstructed in any given sample (i.e., the proportion of samples is 100%). These lncRNAs could only be assembled after pooling data from multiple samples. Data for this figure is given in Table S22.

In many cases, full-length sheep lncRNAs cannot be reconstructed using all reads sequenced from a given individual. For instance, the known lncRNA ENSOARG00000025201 is reconstructed by 28 of the 63 possible GTFs, but none of these GTFs was built using reads from only one individual (Table S21). Only 189 lncRNAs (1.5%) were fully reconstructed in all 63 possible GTFs. Notably, 154 of these are known Ensembl lncRNAs (Table S21).

### lncRNAs are enriched in the vicinity of co-expressed protein-coding genes

Enhancer sequences positively modulate the transcription of nearby genes (see reviews [75, 76]), and may be the evolutionary origin of a fraction of these lncRNAs (as suggested by [77, 78]), including a novel class of enhancer-transcribed ncRNAs, enhancer (eRNAs), which – although a distinct subset – are arbitrarily classified as lncRNAs [79]. eRNAs are likely to be co-expressed with protein-coding genes in their immediate genomic vicinity.

To identify co-regulated sets of protein-coding and non-coding loci, we performed network cluster analysis of the sheep expression level dataset (Table S19) using the Markov clustering (MCL) algorithm [80], as implemented by Graphia Professional (Kajeka Ltd., Edinburgh, UK) (see Methods) [81, 82]. To reduce noise, only those novel lncRNAs with reproducible expression (that is, having > 0.01 TPM in every replicate of the tissue in which it is most highly expressed) are included in this analysis (n = 5353). The resulting graph contained only genes with tightly correlated expression profiles (Pearson’s *r* ≥ 0.95) (Figure 4) and was highly structured, organised into clusters of genes with a tissue or cell-type specific expression profile (Table S23).

**Figure 4.**
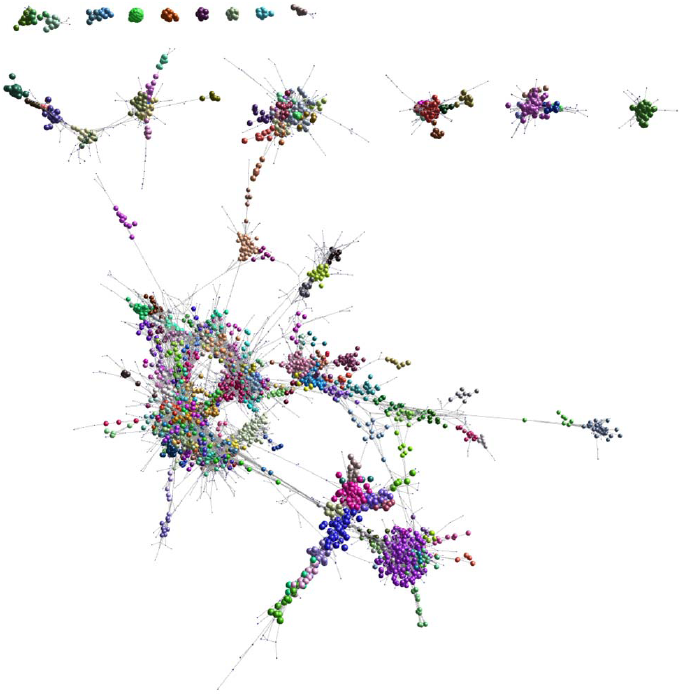
3D visualisation of a gene-to-gene correlation graph. Each node (sphere) represents a gene. Nodes are connected by edges (lines) that represent Pearson’s correlations between the two sets of expression level estimates, at a threshold greater than or equal to 0.95. The graph comprises 11,841 nodes and 2,214,099 edges. Genes cluster together according to the similarity of their expression profiles (i.e. their degree of co-expression), with clusters (coloured sets of nodes) determined using the MCL algorithm. Expression level estimates for the lncRNAs in this graph are given in Table S19. The genes comprising each co-expression cluster are given in Table S23. Those lncRNAs co-regulated with protein-coding genes will be found within the same co-expression cluster.

We expect that for a given cluster of co-expressed genes (which contains *x* lncRNAs and *y* protein-coding genes, each on chromosome *z*), the distance between an enhancer-derived lncRNA and the nearest protein-coding gene should be significantly shorter than the distance between that lncRNA and a random subset of protein-coding genes. For the purposes of this test, each random subset, of size *y*, is drawn from the complete set of protein-coding genes on the same chromosome *z* (that is, the same chromosome as the lncRNA), irrespective of strand and their degree of co-expression with the lncRNA. The significance of any difference in distance was then assessed using a randomisation test (see Methods).

Of the 5353 lncRNAs included in the analysis, 1351 (25%) were found on the same chromosome as a highly co-expressed protein-coding gene (Table S24), with 252 of these (19%) significantly closer to the co-expressed gene than to randomly selected genes from the same chromosome (p < 0.05; Table S25).

Even where the lncRNA is reproducibly expressed in each of 6 animals, there is still substantial noise in the expression estimates with compromises co-expression analysis. We therefore calculated the Pearson’s *r* between the expression profile of each reproducibly expressed lncRNA and its nearest protein-coding gene (which may overlap it), located both 5’ and 3’ on the sheep genome (Table S26). The distance to the nearest gene correlates negatively with the absolute value of Pearson’s *r*, both for genes upstream (*rho* = -0.19, p < 2.2x10^−16^) and downstream (*rho* = -0.21, p < 2.2x10^−16^) of the lncRNA (Table S26). This suggests that, in general, the expression profile of a lncRNA is more similar to nearer than more distant protein-coding genes. Using a variant of the above randomisation test, we also tested whether the absolute value of Pearson’s *r*, when correlating the expression profiles of the lncRNA and its nearest protein-coding gene, was significantly greater than the value of *r* obtained when correlating the lncRNA with 1000 random protein-coding genes drawn from the same chromosome. For this test, analysis was restricted to those lncRNAs on complete chromosomes rather than the smaller unplaced scaffolds. 27% of lncRNA had a Pearson correlation of > 0.5 with either the nearest upstream or downstream gene, and in around 20% of cases, correlation was significantly different (p < 0.05) from the average correlation with the random set (Table S26).

## Conclusion

Comparative analysis of lncRNAs assembled using RNA-seq data from several closely related species – sheep, goat and cattle – demonstrates that for the *de novo* assembly of lncRNAs requires very high-depth RNA-seq datasets with a large number of replicates (> 6 replicates per sample, each sequencing >> 100 million reads). The transcription of many lncRNAs identified by this cross-species approach is conserved, effectively validating their existence. We identified a subset of lncRNAs in close proximity to protein-coding genes with which they are strongly co-expressed, consistent with the evolutionary origin of some ncRNAs in enhancer sequences. Conversely, the majority of lncRNA do not share transcriptional regulation with neighbouring protein-coding genes. Overall, alongside substantially expanding the lncRNA repertoire for several livestock species, we demonstrate that the conventional approach to lncRNA detection – that is, species-specific *de novo* assembly – can be reliably supplemented by data from related species.

## Materials and Methods

### Sheep RNA-sequencing data

We have previously created an expression atlas for the domestic sheep [45], using RNA-seq data largely collected from adult Texel x Scottish Blackface (TxBF) sheep. Experimental protocols for tissue collection, cell isolation, RNA extraction, library preparation, RNA sequencing and quality control are as previously described [45], and independently available on the FAANG Consortium website (http://ftp.faang.ebi.ac.uk/ftp/protocols). All RNA-seq libraries were prepared by Edinburgh Genomics (Edinburgh Genomics, Edinburgh, UK) and sequenced using the Illumina HiSeq 2500 sequencing platform (Illumina, San Diego, USA).

The majority of these libraries were sequenced to a depth of >25 million paired-end reads per sample using the Illumina TruSeq mRNA library preparation protocol (polyA-selected) (Illumina; Part: 15031047, Revision E). A subset of 11 transcriptionally rich ‘core’ tissues (bicep muscle, hippocampus, ileum, kidney medulla, left ventricle, liver, ovary, reticulum, spleen, testes, thymus), plus one cell type in two conditions (bone marrow derived macrophages (BMDMs), unstimulated and 7 hours after simulation with lipopolysaccharide (LPS)), were sequenced to a depth of >100 million paired-end reads per sample using the Illumina TruSeq total RNA library preparation protocol (rRNA-depleted) (Illumina; Part: 15031048, Revision E).

Sample metadata for all tissue and cell samples are deposited in the EBI BioSamples database under submission identifier GSB-718 (https://www.ebi.ac.uk/biosamples/groups/SAMEG317052). The raw read data, as .fastq files, are deposited in the European Nucleotide Archive (ENA) under study accession PRJEB19199 (http://www.ebi.ac.uk/ena/data/view/PRJEB19199).

### Goat RNA-sequencing data

All RNA-seq libraries for goat were prepared by Edinburgh Genomics (Edinburgh Genomics, Edinburgh, UK) (as above) and sequenced using the Illumina HiSeq 4000 sequencing platform (Illumina, San Diego, USA). These libraries were sequenced to a depth of >30 million paired-end reads per sample using the Illumina TruSeq mRNA library preparation protocol (polyA-selected) (Illumina; Part: 15031047, Revision E). Sample metadata for all tissue and cell samples are deposited in the EBI BioSamples database under submission identifier GSB-2131 (https://www.ebi.ac.uk/biosamples/groups/SAMEG330351). The raw read data, as .fastq files, are deposited in the ENA under study accession PRJEB23196 (http://www.ebi.ac.uk/ena/data/view/PRJEB23196).

### Identifying candidate lncRNAs in sheep and goats

We have previously described an RNA-seq processing pipeline for sheep [45] – using the HISAT2 aligner [50] and StringTie assembler [51] – for generating a uniform, non-redundant set of *de novo* assembled transcripts. The same pipeline is applied to the goat RNA-seq data. This pipeline culminates in a single file per species, merged.gtf; that is, the output of StringTie ‐‐merge, which collates every transcript model from the 54 goat assemblies (each assembly being both individual‐ and tissue-specific), and 429 of the 441 assemblies within the sheep expression atlas [45] (12 sheep libraries were not used for this purpose as they were replicates of pre-existing bone marrow-derived macrophage libraries, prepared using an mRNA-seq rather than a total RNA-seq protocol). Not all transcript models in either GTF will be stranded. This is because HISAT2 infers the transcription strand of a given transcript by reference to its splice sites; this is not possible for single exon transcripts, which are unspliced.

The GTF was parsed to distinguish candidate lncRNAs from assembly artefacts, and from other RNAs, by applying the filter criteria of Ilott, *et al.* [52], excluding gene models that (a) were < 200bp in length, (b) overlapped (by >= 1bp on the same strand) any coordinates annotated as ‘protein-coding’ or ‘pseudogene’ (this classifications are explicitly stated in the Ensembl-hosted Oar v3.1 annotation and assumed true of all gene models in the ARS1 annotation), or (c) were associated with multiple transcript models (which are more likely to be spurious). For single-exon gene models, we used a more conservative length threshold of 500bp – the lower threshold of 200bp could otherwise be met by a single pair of reads. We further excluded any novel gene model that was previously considered protein-coding in each species’ expression atlas (as described in [45]); these models contain an ORF encoding a peptide homologous to a ruminant protein in the NCBI nr database [45]. These criteria establish longlists of 30,677 candidate sheep lncRNAs (14,862 of which are multi-exonic) and 7671 candidate goat lncRNAs (3289 of which are multi-exonic). The sheep genome, Oar v3.1, already contains 1858 lncRNA models, of which the StringTie assembly precisely reconstructs 1402 (75%). Despite this pre-existing support, these models were included on the sheep longlist for independent verification. The goat genome, by contrast, was annotated with a focus on protein-coding gene models [83], by consolidating protein and cDNA alignments – from exonerate [84] and tblastn [56] – with the annotation tool EVidence Modeller (EVM) [85]. Consequently, there are no unambiguous lncRNAs in the associated GTF (ftp://ftp.ncbi.nlm.nih.gov/genomes/all/GCF/001/704/415/GCF_001704415.1_ARS1/GCF_001704415.1_ARS1_genomic.gff.gz, accessed 23^rd^ October 2017) (unlike the Ensembl-hosted sheep annotation, the goat annotation is currently only available via NCBI).

Each longlist of candidates was assessed for coding potential using three different tools: CPAT v1.2.3 [54], which assigns coding probabilities to a given sequence based on differential hexamer usage [86] and Fickett TESTCODE score [87], PLEK v1.2, a support vector machine classifier utilising k-mer frequencies [55], and CPC v0.9-r2 [53], which was used in conjunction with the non-redundant sequence database, UniRef90 (the Uniref Reference Cluster, a clustered set of sequences from the UniProt KnowledgeBase that constitutes comprehensive coverage of sequence space at a resolution of 90% identity) [88,89] (ftp://ftp.uniprot.org/pub/databases/uniprot/uniref/uniref90/uniref90.fasta.gz, accessed 18^th^ August 2017). CPC scores putatively coding sequences positively and non-coding sequences negatively. We retained only those sequences with a CPC score < -0.5 (consistent with previous studies [31, 90]) and a CPAT probability < 0.58 (after creating sheep-specific coding and non-coding CPAT training data, from Oar v3.1 CDS and ncRNA, this cut-off is the intersection of two receiver operating characteristic curves, obtained using the R package ROCR [91]; this cut-off is also used for the goat data, as there are insufficient non-coding training data for this species).

For each remaining gene model, we concatenated its exon sequence and identified the longest ORF within it. Should CPC, CPAT or PLEK make a false positive classification of ‘non-coding’, this translated ORF was considered the most likely peptide encoded by the gene. Gene models were further excluded if the translated ORF (a) contained a protein domain, based on a search by HMMER v3.1b2 [57] of the Pfam database of protein families, v31.0 [60], with a threshold E-value of 1x10^−5^, or (b) shared homology with a known peptide in the Swiss-Prot March 2016 release [58, 59], based on a search with BLAST+ v2.3.0 [56]: blastp with a threshold E-value of 1x10^−5^. Shortlists of 12,296 (sheep) and 2657 (goat) candidate lncRNAs – each with three independent ‘non-coding’ classifications and no detectable blastp and HMMER hits – are given in Tables S1 and S2, respectively.

### Classification of lncRNAs

Using the set of Oar v3.1 transcription start sites (TSS), obtained from Ensembl BioMart [92], and the set of ARS1 gene start sites (ftp://ftp.ncbi.nlm.nih.gov/genomes/all/GCF/001/704/415/GCF_001704415.1_ARS1/GCF_001704415.1_ARS1_genomic.gff.gz, accessed 23^rd^ October 2017), we classified novel candidate lncRNAs for each species in the manner of [93], as either (a) sense or antisense (if the coordinates of the lncRNA overlap, or are encapsulated by, a known gene on the same, or opposite, strand), (b) up‐ or downstream, and on the same or opposite strand (if < 5kb from the nearest TSS), or (c) intergenic (if ≥ 5kb, 10kb, 20kb, 50kb, 100kb, 500kb or 1 Mb from the nearest TSS, irrespective of strand). The HISAT2/StringTie pipeline, used to generate these transcript models, cannot infer the transcription strand in all cases, particularly for single-exon transcripts. Accordingly, some lncRNAs will overlap the coordinates of a known gene, but its strandedness with respect to that gene – whether it is sense or antisense – will be unknown.

### Conservation of lncRNAs in terms of sequence

To assess the sequence-level conservation of sheep and goat lncRNA transcripts, we obtained human lncRNA sequences from two databases, NONCODE v5 [62] (http://www.noncode.org/datadownload/NONCODEv5_human.fa.gz, accessed 27^th^ September 2017) and lncRNAdb v2.0 [63] (http://www.lncrnadb.com/media/cms_page_media/10651/Sequences_lncrnadb_27Jan2015.csv, accessed 27^th^ September 2017) (which contain 172,216 and 152 lncRNAs, respectively). A previous study of lncRNAs in cattle [31] also generated a conservative set of 9778 lncRNAs, all of which were detectably expressed in at least one of 18 tissues (read count > 25 in each of three replicates per tissue). These sets of sequences constitute three independent BLAST databases. For each sheep and goat lncRNA, blastn searches [56] were made against each database using an arbitrarily high E-value of 10, as substantial sequence-level conservation was not expected.

### Conservation of lncRNAs in terms of synteny

For each of the human (GRCh38.p10), sheep (Oar v3.1), cattle (UMD3.1) and goat (ARS1) reference genomes, we established those regions in each pairwise comparison where gene order is conserved, obtaining reference annotations from Ensembl BioMart v90 [92] (sheep, cattle and human) and NCBI (goat; ftp://ftp.ncbi.nlm.nih.gov/genomes/all/GCF/001/704/415/GCF_001704415.1_ARS1/GCF_001704415.1_ARS1_genomic.gff.gz, accessed 27^th^ September 2017). By advancing a sliding window across each chromosome gene-by-gene from the 5’ end, we identified the first upstream and first downstream gene of each focal gene, irrespective of strand. For the purpose of this analysis, the first and last genes on each chromosome are excluded, having no upstream or downstream neighbour, respectively. For each pairwise species comparison, we then determined which set of blocks were present in both – that is, where the HGNC symbols for upstream gene/focal gene/downstream gene were identical. These syntenic blocks, of three consecutive genes each, are regions in the genome where gene order is conserved both up‐ and downstream of a focal gene: between sheep and cattle, there are 2927 regions (comprising 5601 unique genes); sheep and goat, 2038 regions (3883 unique genes); cattle and goat, 2982 regions (5258 unique genes); sheep and human, 380 regions (930 unique genes); goat and human, 527 regions (1262 unique genes); cattle and human, 443 regions (1063 unique genes). If in each syntenic block a lncRNA was found between the upstream and focal gene, or the focal and downstream gene, in only one of the two species, a global alignment was made between the transcript and the intergenic region of the corresponding species. Alignments were made using the Needleman-Wunsch algorithm, as implemented by the ‘needle’ module of EMBOSS v6.6.0 [94], with default parameters. By effectively treating lncRNA transcripts as if they were CAGE tags (that is, short reads of 20-50 nucleotides [95]), we considered successful alignments to be those containing one or more consecutive runs of 20 identical residues, without gaps (the majority of these alignments in any case have >= 75% identity across the entire length of the transcript (Table S18)). The probability that a transcript randomly matches 20 consecutive residues, within a pre-defined region, is extremely low.

For successful alignments, the target sequence (that is, an extract from the intergenic region) was considered a novel lncRNA. For this analysis, the sheep and goat lncRNAs used are those from their respective shortlists (Tables S1 and S2). lncRNA locations in other species are obtained from previous studies applying similarly conservative classification criteria. For cattle, 9778 lncRNAs were obtained [31], each of which were >200bp, considered non-coding by the classification tools CPC [53] and CNCI [96], lacked sequence similarity to the NCBI nr [45] and Pfam databases [60], and had a normalised read count > 25 in at least 2 of 3 replicates per tissue for 18 tissues. For human, 17,134 lncRNAs were obtained [72], each of which were assembled from >250bp transfrags, considered non-coding by the classification tool CPAT [54], lacked sequence similarity to the Pfam database [60], and had active transcription confirmed by intersecting intervals surrounding the transcriptional start site with chromatin immunoprecipitation and sequencing (ChIP-seq) data from 13 cell lines.

### Expression level quantification

For the 11 ‘core’ tissues of the sheep expression atlas, plus unstimulated and LPS-stimulated BMDMs (detailed in S2 Table of [45] and available under ENA accession PRJEB19199), expression was quantified using Kallisto v0.43.0 [66] with a k-mer index (k=31) derived after supplementing the Oar v3.1 reference transcriptome with the shortlist of 11,646 novel sheep lncRNA models (Table S1) and those lncRNAs assembled in either human (n = 18), goat (n=164), or cattle (n=1219), and which map to a conserved region of the sheep genome (Table S15). Oar v3.1 transcripts were obtained from Ensembl v90 [92] in the form of separate files for 22,823 CDS (ftp://ftp.ensembl.org/pub/release-90/fasta/ovis_aries/cds/Ovis_aries.Oar_v3.1.cds.all.fa.gz, accessed 27^th^ September 2017) and 6005 ncRNAs (ftp://ftp.ensembl.org/pub/release-90/fasta/ovis_aries/ncrna/Ovis_aries.Oar_v3.1.ncrna.fa.gz, accessed 27^th^ September 2017). An equivalent set of expression estimates was made for goat, across the 21 tissues and cell types of the goat expression atlas (i.e., 54 RNA-seq libraries available under ENA accession PRJEB23196). 47,193 transcripts, from assembly ARS1, were obtained from NCBI
(ftp://ftp.ncbi.nlm.nih.gov/genomes/all/GCF/001/704/415/GCF_001704415.1_ARS1/GCF_001704415.1_ARS1_rna.fna.gz, accessed 27^th^ September 2017), and supplemented both with the shortlist of 2657 novel goat lncRNA models (Table S2), and those lncRNAs assembled in human (n = 15), sheep (n = 507), or cattle (n= 1213) (Table S15). After quantification in each species, transcript-level abundances were summarised to the gene-level.

### Categorisation of expression profiles

Expression levels were categorised in the manner of the Human Protein Atlas [97], and as previously employed in the Sheep Gene Expression Atlas [45]. Each gene is considered to have either no expression (average TPM < 1, a threshold chosen to minimise the influence of stochastic sampling), low expression (10 > average TPM >= 1), medium expression (50 > average TPM > 10), or high expression (average TPM >= 50). Two sample specificity indices were calculated for each gene, as in [45]: firstly, *tau*, a scalar measure of expression breadth bound between 0 (for housekeeping genes) and 1 (for genes expressed in one sample only) [70], and secondly, the mean TPM (across all samples) divided by the median TPM (across all tissues). Genes with greater sample specificity will have a more strongly skewed distribution (i.e. a higher mean and a lower median), and so the larger the ratio, the more sample-specific the expression. To avoid undefined values, should median TPM be 0, it is considered instead to be 0.01.

Each gene is also assigned one or more categories, to allow an at-a-glance overview of its expression profile: (a) ‘tissue enriched’ (expression in one tissue at least five-fold higher than all other tissues [‘tissue specific’ if all other tissues have 0 TPM]), (b) ‘tissue enhanced’ (five-fold higher average TPM in one or more tissues compared to the mean TPM of all tissues with detectable expression [this category is mutually exclusive with ‘tissue enriched’), (c) ‘group enriched’ (five-fold higher average TPM in a group of two or more tissues compared to all other tissues (‘groups’ are analogous to organ systems, and are as described in the sheep expression atlas [45]), (d) mixed expression (detected in one or more tissues and neither of the previous categories), (e) ‘expressed in all’ (>= 1 TPM in all tissues), and (f) ‘not detected’ (< 1 TPM in all tissues).

### Network analysis

Network analysis of the sheep expression level data was performed using Graphia Professional (Kajeka Ltd, Edinburgh, UK), a commercial version of BioLayout *Express*^3D^ [81, 82]. A correlation matrix was built for each gene-to-gene comparison, which was then filtered by removing all correlations below a given threshold (Pearson’s *r* < 0.95). A network graph was then constructed by connecting nodes (genes) with edges (correlations above the threshold). The local structure of the graph – that is, clusters of co-expressed genes (detailed in Table S23) – was interpreted by applying the Markov clustering (MCL) algorithm [80] at an inflation value (which determines cluster granularity) of 2.2.

### Enrichment of lncRNAs in the vicinity of protein-coding genes

To test whether lncRNAs co-expressed with protein-coding genes are more likely to be closer to them (from which we can infer they are more likely to have been derived from an enhancer sequence affecting that protein-coding gene), we employed a randomisation test in the manner of [98]. We first obtained clusters of co-expressed genes from a network graph of the sheep expression level dataset (see above). We then calculated *q*, the number of times the distance between each lncRNA and the nearest protein-coding gene within the same cluster was higher than the distance between each lncRNA and the nearest gene within *s* = 1000 randomly selected, equally sized, subsets of protein-coding genes, drawn from the same chromosome as each lncRNA. Letting *r* = *s*-*q*, then the p-value of this test is *r*+1/*s*+1.

## Declarations

### Acknowledgements

The authors would like to thank the farm staff at Dryden farm and members of the sheep tissue collection team from The Roslin Institute and R(D)SVS who were involved in tissue collections for the sheep gene expression atlas project. Rachel Young and Lucas Lefevre isolated the bone marrow derived macrophages and Zofia Lisowski provided technical assistance with collection and post mortem for the goat samples. Technical expertise for dissection of the sheep brain samples was provided by Fiona Houston and heart samples by Kim Summers and Hiu-Gwen Tsang. The authors are also grateful for the support of the FAANG Data Coordination Centre in the upload and archiving of the sample data and metadata.

### Ethics approval and consent to participate

Approval was obtained from The Roslin Institute’s and the University of Edinburgh’s Protocols and Ethics Committees. All animal work was carried out under the regulations of the Animals (Scientific Procedures) Act 1986.

### Data availability

The raw RNA-sequencing data are deposited in the European Nucleotide Archive (ENA) under study accessions PRJEB19199 (sheep) and PRJEB23196 (goat). Sample metadata for all tissue and cell samples, prepared in accordance with FAANG consortium metadata standards, are deposited in the EBI BioSamples database under group identifiers SAMEG317052 (sheep) and SAMEG330351 (goat). All experimental protocols are available on the FAANG consortium website at http://ftp.faang.ebi.ac.uk/ftp/protocols.

## Supplementary Material

**Dataset S1.** 63 sequence assemblies (as GTFs): all possible combinations when merging 6 different sets of RNA-seq reads (available via the University of Edinburgh DataShare portal; http://dx.doi.org/10.7488/ds/2284).

**Table S1.** Candidate sheep lncRNAs: a shortlist of novel gene models (plus independently confirmed known gene models) assessed for coding potential using CPC, CPAT, PLEK, blastp vs. Swiss-Prot, and HMMER vs. Pfam.

**Table S2.** Candidate goat lncRNAs: a shortlist of novel gene models assessed for coding potential using CPC, CPAT, PLEK, blastp vs. Swiss-Prot, and HMMER vs. Pfam.

**Table S3.** Sheep gene models considered non-coding by either CPC, CPAT or PLEK but showing sequence homology to either a known protein (in Swiss-Prot) or protein domain (in Pfam-A).

**Table S4.** Goat gene models considered non-coding by either CPC, CPAT or PLEK but showing sequence homology to either a known protein (in Swiss-Prot) or protein domain (in Pfam-A).

**Table S5.** Number of novel sheep lncRNA gene models identified per chromosome.

**Table S6.** Number of novel goat lncRNA gene models identified per chromosome.

**Table S7.** Number of novel sheep lncRNA gene models identified, by category.

**Table S8.** Number of novel goat lncRNA gene models identified, by category.

**Table S9.** Alignments of novel sheep lncRNA gene models to goat, cattle and human lncRNAs.

**Table S10.** Alignments of novel goat lncRNA gene models to sheep, cattle and human lncRNAs.

**Table S11.** Presence of intergenic lncRNAs both in sheep and cattle, in regions of conserved synteny.

**Table S12.** Presence of intergenic lncRNAs both in sheep and goat, in regions of conserved synteny.

**Table S13.** Presence of intergenic lncRNAs both in cattle and goat, in regions of conserved synteny.

**Table S14.** Presence of intergenic lncRNAs both in sheep and human, in regions of conserved synteny.

**Table S15.** Presence of intergenic lncRNAs both in goat and human, in regions of conserved synteny.

**Table S16.** Presence of intergenic lncRNAs both in cattle and human, in regions of conserved synteny.

**Table S17.** High-confidence lncRNA pairs, those conserved across species both sequentially and positionally.

**Table S18.** lncRNAs inferred in one species by the genomic alignment of a transcript assembled with the RNA-seq libraries from a related species.

**Table S19.** Expression level estimates for 13,047 novel sheep lncRNAs, as transcripts per million (TPM), assessed using 71 adult RNA-seq libraries (11 tissues plus one cell type in two different conditions, each sequenced in 6 individuals).

**Table S20.** Expression level estimates for 4392 novel goat lncRNAs, as transcripts per million (TPM), assessed using 54 RNA-seq libraries (20 tissues plus one cell type in two different conditions, each sequenced in 4 individuals).

**Table S21.** Reproducibility of sheep lncRNA gene models when merging all combinations of data from 6 adults (3 female, 3 male), each individual having sequenced a common set of RNA-seq libraries (comprising 31 tissues/cell types).

**Table S22.** Number of sheep expression atlas RNA-seq libraries (out of 429 in total) in which a candidate lncRNA gene model cannot be fully reconstructed.

**Table S23.** Genes within each co-expression cluster, after network analysis of the sheep RNA-seq libraries.

**Table S24.** No. of lncRNAs co-expressed with protein-coding genes.

**Table S25.** Distance between lncRNAs and protein-coding genes within the same coexpression cluster, on the same chromosome.

**Table S26.** Correlation between the expression profile of sheep lncRNAs and their nearest protein-coding genes, both 5’ and 3’.

